# Yield and agronomic performance of sweet corn in response to inoculation with *Azospirillum* sp. in arid land conditions

**DOI:** 10.1101/2023.04.27.538588

**Authors:** Sergio Contreras-Liza, Christofer Villadeza, Pedro Rodriguez Grados, Edison Goethe Palomares, Carlos Irwin Arbizu

## Abstract

Nitrogen is the most common limiting factor for crop productivity and most maize cultivars require fertilizing. Here we report the possibility of partially replacing the nitrogenous fertilizer in sweet corn inoculated with a native strain of Azospirillum in arid land on the coast of Peru. We examined an agronomic experiment in a crop field of arid soils under drip irrigation in Huacho (Peru) using a commercial variety of sweet corn. The treatments were two levels of nitrogen (90 and 180 kg N ha^-1^), one or two applications to the foliage of a native strain of *Azospirillum* sp. (1 × 10^8^ CFU/mL) and a control treatment only with nitrogen fertilizer. Eleven agronomic variables related to productive aspects were evaluated, performing statistical analysis and the comparison of treatment means. The inoculation with *Azospirillum* sp. did not significantly (*p*> 0.05) affect the total weight of ears, the number of ears per plant and the number of male flowers, but it significantly (*p*< 0.05) influenced the grain yield per hectare, survival of plants, the weight of grain per plant, diameter and length of the cob. In some productive characteristics of sweet corn cv“Pardo”, a significant effect was found by inoculation with *Azospirillum* sp., surpassing in grain yield the control only with nitrogen fertilization, suggesting that it is possible to complement the application of nitrogen to the soil through the inoculation of this strain, replacing up to 50% of the levels of fertilizer application since the B/C ratio increased.

## Introduction

Corn (*Zea mays*) is one of the most important agricultural products in Latin America; in this region, local maize varieties are critical for food security, livelihoods, and cultures (Guzzon et al. 2021). In Peru, the area under cultivation exceeds half a million hectares (MINAGRI 2017) and the local variety “choclo” (sweet corn) is a highly demanded food in the market of its importance in regional gastronomy, as Andean maize is the basis of the rural population (Salvador-Reyes and Pedrosa 2020). Due to the need to obtain sustainable agricultural production, new ecological alternatives have been investigated worldwide, considering the use of biofertilizers as an option to partially or totally replace the use of chemical fertilizers (Shovitri et al. 2022). Recent studies reinforce the concept that corn yields are controlled not only by the N supply in the soil but also by factors that modify the demand of the plants and the ability to capture N (Correndo et al. 2021).

The arid soils of the Peruvian coast have a low amount of organic matter, normally less than 1 % (Yang 2020); the balance between mineralization and decomposition of soil organic matter is due to functional differences in key soil microbial groups that influence the mineralization of carbon sources (Whitaker et al. 2014). For this reason, it is necessary to explore those microbial groups that would be related to the biological fixation of atmospheric nitrogen and the mineralization of soil organic matter, which have an effect on the agronomic performance of maize (Shovitri et al. 2022). Likewise, corn producers need to improve crop management, especially in technological aspects of nitrogen fertilization that reduce high production costs (Morris et al. 2018).

Azospirillum develops in the roots supplying N to the plant and in various edaphic environments associated with cereals and other species; they are capable of synthesizing phytohormones that will promote growth and morphological and physiological changes of the root and exert biocontrol (Correa et al. 2007), improving the use of water and nutrients and increasing yield and productivity (Dobbelaere et al. 2001). Azospirillum was originally selected for its ability to fix atmospheric nitrogen (N_2_). Since the 1970s, it has consistently shown to be a very promising PGPR, based on physiological, molecular, agricultural, and environmental studies conducted with this bacteria (Bashan et al. 2013). In recent years, studies on Azospirillum-plant interactions have introduced a wide range of mechanisms to demonstrate the beneficial impacts of this bacterium on plant growth (Cassan et al. 2020); alternative strategies have been developed to replace chemical fertilizers through the use of beneficial microorganisms that maintain soil balance and support plant growth through various mechanisms, including phosphate solubilization and nitrogen fixation (Corrales et al. 2014). The use of plant growth-promoting bacteria (PGPR) for the formulation of bio fertilisers has become one of the most promising clean technologies for the development of sustainable agriculture (Vejan et al. 2016). Among these bacteria, the ones that stand out the most is Azospirillum, which has the ability to fix nitrogen, produce cytokinins, gibberellins and indols, and reduce nitrates, which allows it to be used as a biofertilizer without generating consequences for the environment (Bashan et al., 2013; Fibach-Paldi et al. 2012). Through direct and indirect mechanisms of action, these bacteria can significantly reduce the use of chemical fertilizers (Adesemoye et al. 2009).

Nitrogen fixation is the main mechanism by which Azospirillum influences plant growth (Fukami *e*t al. 2018). In recent years, some studies have focused on the nitrogen cycle within cells and the genes involved in this process. It appears that the ability of Azospirillum strains is often naturally maintained, enhancing their ability to express exceptional nitrogenase enzyme activity (Goswami *et al*. 2016). The efficiency for atmospheric nitrogen fixation and denitrification can be regulated through the concentration of oxygen, nitrates and molybdenum, in addition, Azospirillum has the ability to adapt to low temperatures and low oxygen concentrations, depending on the ability of the bacteria to efficiently use nitrites and nitrates (Tsagou et al., 2003).

According to Walters et al. (2018), some indigenous varieties of maize grown under traditional agricultural practices with little or no fertilizer, have developed strategies to improve plant performance under conditions of low nitrogen content in the soil and in those varieties of maize, 29% 82% of the assimilated nitrogen was derived from the atmospheric form N_2_. According to the research of Zambonin et al. (2019), no interaction was found between the maize hybrids and the inoculation with Azospirillum in any variable studied and the specificity between the hybrids and the inoculation was not verified. Likewise, they found that the inoculation with *A. brasilense* did not interfere with the grain yield and the yield components of the corn crop; the use of this diazotrophic bacteria seems to be viable even when high doses of N are applied (Galindo et al. 2016).

Rangel-Lucio et al. (2011) found that there is some degree of affinity or effect of the homologous strain between Azospirillum obtained from H-28 and Chalqueño maize and its inoculation in these varieties when evaluating the number of azospirilas and their nitrogenases, as well as a recognition of Azospirillum strains from corn of recent origin, due to the ancestors of this cereal. Likewise, in another investigation carried out by Rangel-Lucio *et al*. (2014), it was found that the biofertilization of *Azospirillum* spp. showed potential benefits in the production of sorghum grain, and in particular, the strains of *A. brasilense* VS-7 and VS-9, presenting an effect of 55% and 49% higher than the control fertilized with nitrogen; their results also demonstrated the affinity between the bacterial strain and the plant genotype.

Sangoquiza et al. (2018) investigated the biological response of *Azospirillum* spp. against different types of stress, for which they carried out the morphological characterization of the isolates as well as their biological response to stress conditions of temperature, salinity, and pH. Their results showed that freeze-dried isolates of *Azospirillum* spp. grow best at temperatures between 28°C and 38°C and pH between 7 and 8; in relation to salinity, some isolates showed good growth in up to 3.5% NaCl.

The objective of this study was to determine the influence of inoculation with a native strain of *Azospirillum* sp., in addition to nitrogen fertilization in sweet corn, and to explore a reduction in the use of chemical fertilizers in this crop in arid conditions of the coast of Peru.

## Materials and Methods

### Study Area

The work was carried out in an aridisol (residual formation soil) during the spring of 2019 in “El Paraíso”, province of Huaura, Lima (coordinates -11.188352, -77.492155). The soil presented a loamy sand texture, alkaline reaction (pH 8.1), very low organic matter content (0.36%) and moderate to high salinity (6.46 dS/m).

According to the Köppen climate classification, the study area is defined as BWh (dry, desert and hot climate); climatic records showed relative humidity between 65 and 91%, a maximum temperature of 19°C and a minimum of 13°C, with no rainfall.

### Materials

- Bacterial inoculum: The bacterial strain was isolated from the rhizosphere of maize in fields on the central coast of Peru and identified as *Azospirillum* sp. by the Laboratorio de Biotecnología de la Producción (Universidad Nacional José Faustino Sánchez Carrión), through morphological characterization and biochemical profile (catalase + test, urease + reaction, motility test and oxidase + test). To formulate the biofertilizer based on *Azospirillum* sp., the biomass in nutrient broth was increased and quantified in Colony Forming Units (CFU) obtaining a final concentration of 1x10^8^ CFU/mL
- Plant material: seeds of sweet corn variety. “Pardo”, was purchased from a local seed supplier.

### Treatment application procedures

- Inoculation: A first dose of 30 mL of inoculum of *Azospirillum* sp. (1x10^8^ CFU) as a seed treatment was applied. This dose was applied according to the following combinations of nitrogen fertilization and inoculants: T_0_ control (without inoculation) 180 kg N ha^-1^, T_1_ one application of Azospirillum in the seed + 180 kg N ha^-1^, T_2_ two applications of Azospirillum (in the seed and at hilling) +180 kg N ha^-1^, T_3_ one application of Azospirillum in the seed + 90 kg N ha^-1^, T_4_ two applications of Azospirillum (in the seed and at hilling) + 90 kg N ha^-1^ (Table 1).

**Table 1.** Description of treatments

| Treatments | Code | Description |
| --- | --- | --- |
| 100%N | Control | Conventional fertilization 180 kg N ha<sup>-1</sup>, uninoculated |
| 100%N + A<sub>1</sub> | T<sub>1</sub> | 180 kg N+ Azospirillum inoculation at sowing |
| 100%N + A<sub>1</sub> + A<sub>2</sub> | T<sub>2</sub> | 180 kg N+ Azospirillum inoculation at sowing and hilling |
| 50%N + A<sub>1</sub> | T<sub>3</sub> | 90 kg N+ Azospirillum inoculation at sowing |
| 50%N + A<sub>1</sub> + A<sub>2</sub> | T<sub>4</sub> | 90 kg N+ Azospirillum inoculation at sowing and hilling |

Nitrogen fertilization treatments were formulated and coded according to Table 2, using the urea source (46% N) and applied 15 and 45 days after planting, according to each level considered. The control treatment (without inoculation) was applied with the full dose of 180 kilos of nitrogen per hectare, dividing the fertilizer at 15 and 45 days after sowing. Treatments T_2_ and T_4_ received the second dose of the inoculum sprayed on the foliage at the time of hilling (45 days after sowing). The plots without bacterial inoculants (T_0_) were only sprayed with running water on the foliage of the plant. All experimental units (inoculated or non-inoculated) received a background application with 100 units of phosphorus and 120 units of potassium per hectare as standard, to complete the fertilization formula. Irrigation was carried out by pumping through drip irrigation tapes to guarantee an adequate water supply to the plants, due to the aridity and soil salinity conditions.

**Table 2.** ANOVA for yield parameters

| Sources | dF | Mean Squares |  |  |  |  |  |
| --- | --- | --- | --- | --- | --- | --- | --- |
|  |  | Yield<br>kg ha<sup>-1</sup> | Grain<br>Weight | Ear<br>Diameter | Ear<br>Length | Ear<br>Weight | Ear<br>Number |
| Blocks | 3 | 459343.25 | 23978.33 | 0.01 | 1.04 | 3.33E-03 | 2.60E-03 |
| Treatments | 4 | **354621.93\*** | 6661.25 | **0.09\*** | **5.64\*** | 7.50E-03 | 1.50E-03 |
| Error | 12 | 68572.15 | 3222.08 | 0.02 | 1.20 | 3.33E-03 | 9.00E-04 |
| CV % |  | 7.70 | 10.06 | 3.00 | 4.97 | 18.35 | 9.77 |
| Means |  | 3400.81 | 564.25 | 4.71 | 22.04 | 0.32 | 1.02 |

Values in **bold\*** are significant at $p < 0.05$

### Evaluated variables

The variables related to maize production that were submitted for evaluation were: yield per hectare (kg), the weight of grain per plot (kg), weight of 100 seeds (kg), cob length (cm), number of cobs per plant and ear diameter (cm). Likewise, days to germination, plant survival to harvest (%), harvest date (days), and the number of spikes per plot. For this, random samples of 10 plants were taken for each variable, except for grain yield, grain weight per plot, days to germination and plant survival, in which the two central rows of each experimental unit were evaluated.

### Analysis of data

A Randomized Complete Block Design with 4 repetitions was used. The size of the experimental unit was 120 maize plants for each treatment; the experimental units were randomly assigned for each inoculation treatment and had a dimension of 4 rows of 10 meters long; the distance between rows was 0.90 m and between plants 0.15 m.

For the hypothesis tests, the level of statistical significance α = 5% was used. Analysis of variance (ANOVA) was performed for the agronomic characteristics evaluated, previously they were subjected to statistical tests to verify the normality of the data, then compared the means of treatments by the Scott-Knott test at a level of significance α= 5%. The data was processed using the R studio program.

## Results

In Table 2, statistical significance can be observed for the variables of yield per hectare, diameter and length of the ear. Still, no differences were found for the characteristics of grain weight per plant, the total weight of the ear and the number of ears.

Likewise, statistical differences were presented for plant survival, days to harvest and weight of 100 seeds. Still, no significance was observed for days to emergence and the number of spikes per plant (Table 3).

**Table 3.** ANOVA for agronomic traits

| Sources | dF | Mean Squares |  |  |  |  |
| --- | --- | --- | --- | --- | --- | --- |
|  |  | Days to<br>emergence | Plant<br>Survival | Harvest Date | Spike<br>number | Seed<br>Weight |
| Blocks | 3 | 1.24 | 211.0 | 137.92 | 14.73 | 3,00E-05 |
| Treatments | 4 | 1.18 | **162.0\*** | **147.2\*** | 58.45 | **9,50E-03\*** |
| Error | 12 | 0.37 | 31.0 | 37.83 | 37.82 | 2,50E-03 |
| CV % |  | 8.95 | 7.64 | 14.09 | 19.28 | 0.54 |
| Means |  | 6.80 | 0.73 | 43.65 | 31.90 | 0.92 |

dF degrees of freedom, CV % coefficient of variability expressed in percentage

Regarding the grain yield per hectare, the treatments with one or two inoculations and with doses of 50 to 100% of the nitrogen level, presented mean values higher than the control without inoculation. On the other hand, for the cob diameter, the treatments with one or two doses of Azospirillum statistically surpassed the control without inoculation, while for the cob length, only the treatment with two inoculations surpassed the control; There were no significant differences for the grain weight per plant and for the number of ears per plant due to the effect of the inoculation treatments compared to the control (Table 4).

**Table 4.** Effect of inoculation + fertilization treatments on yield parameters

| Treatments | Yield<br>kg ha<sup>-1</sup> | Grain<br>Weight, g | Ear<br>Diameter, cm | Ear Length,<br>cm | Ear weight,<br>kg | Ear<br>Number |
| --- | --- | --- | --- | --- | --- | --- |
| 50%N +A<sub>1</sub> | 3633.26<sup>a</sup> | 601.25<sup>a</sup> | 5.06<sup>b</sup> | 21.73<sup>b</sup> | 0.28<sup>b</sup> | 0.92<sup>a</sup> |
| 50%N +A<sub>1</sub>+A<sub>2</sub> | 3269.33<sup>b</sup> | 586.25<sup>a</sup> | 5.12<sup>b</sup> | 20.82<sup>b</sup> | 0.27<sup>b</sup> | 0.96<sup>a</sup> |
| Control (100%N) | 2968.32<sup>b</sup> | 577.50<sup>a</sup> | 5.32<sup>a</sup> | 21.30<sup>b</sup> | 0.35<sup>a</sup> | 1.04<sup>a</sup> |
| 100%N +A<sub>1</sub> | 3709.28<sup>a</sup> | 561.25<sup>a</sup> | 5.39<sup>a</sup> | 23.82<sup>a</sup> | 0.36<sup>a</sup> | 0.95<sup>a</sup> |
| 100%N+A<sub>1</sub>+A<sub>2</sub> | 3434.23<sup>a</sup> | 496.25<sup>a</sup> | 5.37<sup>a</sup> | 22.65<sup>a</sup> | 0.34<sup>a</sup> | 0.88<sup>a</sup> |
| SE | 130.93 | 28.38 | 0.08 | 0.55 | 0.03 | 0.05 |

SE, standard deviation

The effects of inoculation treatments were significant in the case of plant survival at harvest where it was observed that the inoculation with one or two applications of Azospirillum was superior to the control even with a 50% nitrogen contribution to the soil (Table 5). It was also possible to observe that the harvest date in days from sowing was statistically lower (33.5 days) with the treatment of two inoculations and 50% of the applied nitrogen, with respect to the control and the rest of the treatments. Likewise, a slight increase in the weight of the seed was found with the inoculation of Azospirillum plus 100% of the nitrogen applied to the soil, with respect to the non-inoculated control. There were no statistical differences for days to germination and the number of spikes per experimental unit due to the effect of any of the treatments evaluated.

**Table 5.** Effect of inoculation + fertilization treatments on agronomic traits

| Treatments | Days to<br>emergence | Plant<br>Survival % | Harvest<br>Date, days | Flower<br>number | Seed<br>Weight, kg |
| --- | --- | --- | --- | --- | --- |
| 50%N +A<sub>1</sub> | 6.91<sup>a</sup> | 0.78<sup>a</sup> | 43.00<sup>a</sup> | 34.25<sup>a</sup> | 0.93<sup>b</sup> |
| 50%N +A<sub>1</sub>+A<sub>2</sub> | 6.65<sup>a</sup> | 0.70<sup>b</sup> | 33.50<sup>b</sup> | 25.75<sup>a</sup> | 0.94<sup>a</sup> |
| Control (100%N) | 7.50<sup>a</sup> | 0.64<sup>b</sup> | 47.75<sup>a</sup> | 33.50<sup>a</sup> | 0.93<sup>b</sup> |
| 100%N +A<sub>1</sub> | 6.03<sup>a</sup> | 0.79<sup>a</sup> | 48.50<sup>a</sup> | 30.75<sup>a</sup> | 0.94<sup>a</sup> |
| 100%N+A<sub>1</sub>+A<sub>2</sub> | 7.04<sup>a</sup> | 0.74<sup>a</sup> | 45.50<sup>a</sup> | 35.25<sup>a</sup> | 0.93<sup>b</sup> |
| SE | 0.31 | 0.03 | 3.08 | 3.07 | 0.01 |

A<sub>1</sub> first inoculation at sowing, A<sub>2</sub> second inoculation at hilling

Table 6 shows that the effect of inoculation with *Azospirillum* sp. impacts the cost-benefit ratio (B/C) in the production of sweet corn, increasing this ratio from 0.32 to 0.49 when nitrogen levels are reduced to 50% and from 0.23 to 0.35 when applying the full dose of nitrogen fertiliser (180 kg N ha^-1^); in both cases, the effect of inoculation with *Azospirillum* sp. improves the B/C ratio compared to the non-inoculated control.

**Table 6.** Grain yield and benefit-cost ratio (B/C) per hectare (US$)

| Code | Treatment | Yield<sup>1</sup> | GVP<sup>2</sup> | PCost<sup>3</sup> | Benefit/ha | B/C<sup>4</sup> |
| --- | --- | --- | --- | --- | --- | --- |
| T<sub>3</sub> | 50%N +A<sub>1</sub> | 3633.26 | 1963.92 | 1321.08 | 642.84 | 0.49 |
| T<sub>4</sub> | 50%N +A<sub>1</sub>+A<sub>2</sub> | 3269.33 | 1767.21 | 1334.59 | 432.61 | 0.32 |
| T<sub>0</sub> | Control | 2968.32 | 1604.50 | 1469.73 | 134.77 | 0.09 |
| T<sub>1</sub> | 100%N +A<sub>1</sub> | 3709.28 | 2005.02 | 1483.24 | 521.77 | 0.35 |
| T<sub>2</sub> | 100%N+A<sub>1</sub>+A<sub>2</sub> | 3434.23 | 1856.34 | 1510.27 | 346.07 | 0.23 |

<sup>1</sup> Grain yield, kg per ha., <sup>2</sup> Gross value of production, US\$ per ha., <sup>3</sup> Production cost, US\$ per ha.,
<sup>4</sup> Benefit/ha: GVP-PCost (US\$), <sup>5</sup> Benefit/Cost ratio

## Discussion

In the present investigation, it was found that the grain yield in the sweet corn variety “Pardo” was lower with the control dose (180 kg N ha^-1^) without inoculation, compared to the treatments with 90-180 kg N ha^-1^ inoculated with the strain of *Azospirillum* sp. at 15 and 45 days after sowing. This result is consistent with the findings of various authors (Galindo et al. 2020; Schmidt and Gaudin, 2018; Alvarado et al. 2018), which indicate an improvement in corn grain yield under the effect of inoculation with *Azospirillum* sp. The evidence shown by Ferrerira et al. (2020), who found that corn hybrids showed greater expressiveness in yield components in the presence of *A. brasilense* applied in seed treatment, is consistent with this experiment carried out on sweet corn where the seed was also inoculated. Likewise, Alvarado et al. (2018), maintain that the joint use of synthetic fertilizers with microbial inoculants increases grain yield in various varieties of hybrid maize, which shows evidence similar to that obtained in the investigation.

Regarding the other characteristics that are considered yield components evaluated in the experiment, such as ear diameter and length and seed weight, these were significantly affected by inoculation with *Azospirillum* sp., although other variables such as the number of ears, were not affected by inoculation. The significance of the variance components, in this case, suggests that the response in grain yield in this variety of sweet corn can be balanced with nitrogen fertilization and inoculation with *Azospirillum* sp.; this aspect has been studied by Wagner et al. (2020) found that interactions with soil microorganisms are important for the expression of heterosis in corn.

*Azospirillum* spp. and other bacterial strains that have the ability to fix atmospheric nitrogen, solubilize phosphorus or produce growth regulators (Pérez-Montaño et al. 2013; Schmidt and Gaudin, 2018), can be used as biofertilizers in a complementary way to the application of chemical fertilizers (Fukami et al. 2018), a fact that is corroborated in the present investigation under the conditions of the Peruvian coast, in which alluvial soils devoid of organic matter and scarce in water predominate (Yang, 2020).

The intensive use of fertilizers and phytosanitary agents can increase the toxic agents in rivers and soils, and it has been observed that modern intensive agriculture can strongly impact traditional agriculture in desert areas (Lacroix et al. 2020); the environmental impacts are influenced by the high energy intensity linked to the production of inorganic fertilizers used and by phytosanitary agents (Vázquez-Rowe et al. 2016), in this sense, this research is a contribution to evaluate different nitrogen fertilization alternatives in the maize that include the use of soil microbiota such as *Azospirillum* sp. since maize is a product of high demand and deficit in agribusiness (Erenstein et al. 2022). This aspect is especially important in the current world situation of increased fertilizer prices (Brester and Smith, 2022). It is also worth mentioning that Peru and several Latin American countries are importers of fertilizers, due to the absence or deficit in the national production of these inputs; there is an indirect trade effect of international conflicts that can be expected to be more relevant for the economies of Latin America and the Caribbean (Barcena, 2022).

In conclusion, a significant effect was found for inoculation with *Azospirillum* sp. in some productive characteristics of sweet corn cv. “Pardo”, surpassed the control (only with nitrogen fertilization) in grain yield. The grain yields with half the dose of nitrogen applied to the soil (90 kg N/ha) plus an inoculation dose in the sowing of *Azospirillum* sp. was superior to the control treatment without inoculation and with the full dose of nitrogen fertilization (180 kg N/ha). The inoculation with *Azospirillum* sp. improved the survival rate of maize plants in arid soil conditions, devoid of organic matter and with salinity problems. It could be possible to partially replace the nitrogenous fertilizer in the cultivation of sweet corn with *Azospirillum* inoculation since the B/C ratio increases substantially with the use of this bio fertiliser in Peruvian arid soils.

## Acknowledgements

We would like to thank all team members involved in the Laboratorio de Biotecnología de la Producción (Huacho), including Jean Pierre Quiliche for the field sampling and laboratory analysis.

## Author contributions

Sergio Contreras-Liza: data processing and manuscript writing. Christian Yasiel Villadeza: experimental design and field management. Pedro Rodríguez-Grados: strain isolation and characterization. Edison Goethe Palomares: field sampling and evaluation, and Carlos I. Arbizu: manuscript edition.

